# Exploring the mosquito-arbovirus network: a survey of vector competence experiments

**DOI:** 10.1101/2022.05.19.492743

**Authors:** Binqi Chen, Amy R. Sweeny, Velen Yifei Wu, Rebecca Christofferson, Gregory Ebel, Anna C. Fagre, Emily Gallichotte, Rebekah Kading, Sadie J. Ryan, Colin J. Carlson

## Abstract

Arboviruses receive heightened research attention during major outbreaks, or when they cause unusual or severe clinical disease, but are otherwise under-characterized. Global change is also accelerating the emergence and spread of arboviral diseases, leading to time sensitive questions about potential interactions between viruses and novel vectors. Vector competence experiments help determine the susceptibility of certain arthropods to a given arbovirus, but these experiments are often conducted in real-time during outbreaks, rather than with preparedness in mind. We conducted a systematic review of reported mosquito-arbovirus competence experiments, screening 570 abstracts to arrive at 265 studies testing *in vivo* arboviral competence. We found that over 90% of potential mosquito-virus combinations are untested in experimental settings, and that entire regions and their corresponding vectors and viruses are undersampled. These knowledge gaps stymie outbreak response, and limit attempts to both build and validate predictive models of the vector-virus network.

## Introduction

Arthropod-borne (arbo-) viruses face evolutionary pressures that favor generalism in the range of both vertebrate hosts and arthropod vectors that they can use. That flexibility can pose a particular problem for public health, as it both enables their spread into new locations and ecosystems, and adds a layer of unpredictability to their dynamics upon arrival. Experimental studies simplify real-world complexities of transmission, and can be used to test not only the basic compatibility of a given virus and arthropod vector species, but also vector competence—the relative ability of arthropod vectors to be infected by a virus and then disseminate and transmit it to a susceptible host (Mellor, 2000). Despite arboviruses’ evolutionary tendencies towards broad host and vector range (Ciota and Kramer, 2010; Coffey et al., 2008; Kreuder Johnson et al., 2015), there are complex genetic underpinnings that govern vector competence (Beerntsen et al., 2000), which can manifest as variation in competence between closely related species of vector (Kain et al., 2022) or even among populations of the same species (Souza-Neto et al., 2019).

Vector competence experiments are often conducted in response to the emergence of novel pathogens or the emergence of a known pathogen in a new location with previously-untested vectors. The distributions of both mosquito vectors and the viruses they transmit are increasingly in a state of disequilibrium, as a result of climate change, global travel and trade, and biotic homogenization (Kraemer et al., 2019; Ryan et al., 2019). Operating in a responsive paradigm, medical entomology is increasingly struggling to keep pace with these shifts, as resources are often abruptly diverted to new study systems to answer questions that support outbreak response, and influxes of funding to support such studies are typically reactionary to emergence events (Kading et al., 2020). The patchwork of experimental research effort to date represents the cumulative history of these moments, rather than a systematic exploration of the mosquito-arbovirus network, limiting outbreak preparedness, and particularly complicating efforts to predict unrealized links in that network using machine learning (Albery et al., 2021; Evans et al., 2017). As there are no standardized repositories that register vector competence experiments or immortalize their findings, it is currently difficult to evaluate the distribution of research effort so far, and identify important gaps that may be relevant to future outbreaks.

Here, we conducted a systematic review of the mosquito-arbovirus literature using keywords associated with vector competence, and screened 570 studies in Web of Science to identify these experiments. From a total of 265 peer-reviewed studies that met our screening criteria, our objective was to determine the taxonomic and geographic patterns in these studies, and to identify historical trends driving research in this subfield.

## Results

### Taxonomic coverage

We found a total of 298 pairs of viruses (n = 35) and mosquito species (n = 122) that have been tested experimentally, leaving the majority of all possible pairings untested (93%; 3,972 of 4,270 possible pairs; Figure 1). Among these, some viruses were entirely untested in *Aedes* or *Culex* vectors, and most were untested in *Anopheles* (Figure 2). Some untested pairings, such as Everglades virus (EVEV) in *Ae. aegypti*, warrant further investigation given ongoing and future range expansions. However, several pairings for which our search parameters yielded no results in Web of Science were later located in the literature using Google Scholar, such as Murray Valley encephalitis virus (MVEV) and yellow fever (YFV) in *Culex* spp (Davis and Shannon, 1929; Kay et al., 1984, 1982; Kerr, 1932; Whitman et al., 1937), or Ross River virus (RRV) and Sindbis virus (SINV) in *Ae. aegypti* (Hodoameda et al., 2022; Sinclair and Asgari, 2020).

**Figure 1.**
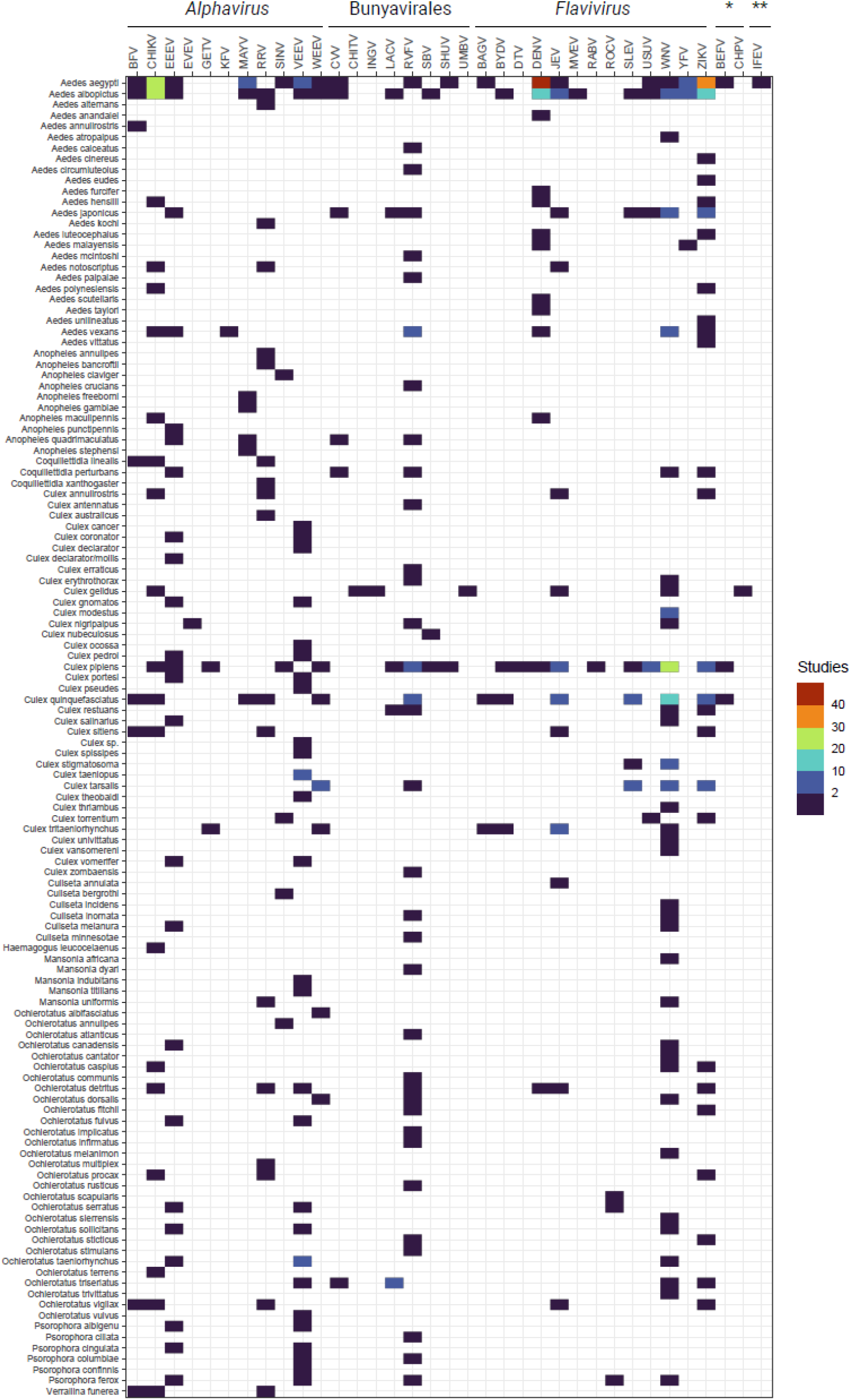
Number of studies that have experimentally tested a given arbovirus-mosquito pair. Abbreviations follow naming conventions in virology. (Key: *Rhabdoviridae; \*\**Orbivirus*)

**Figure 2.**
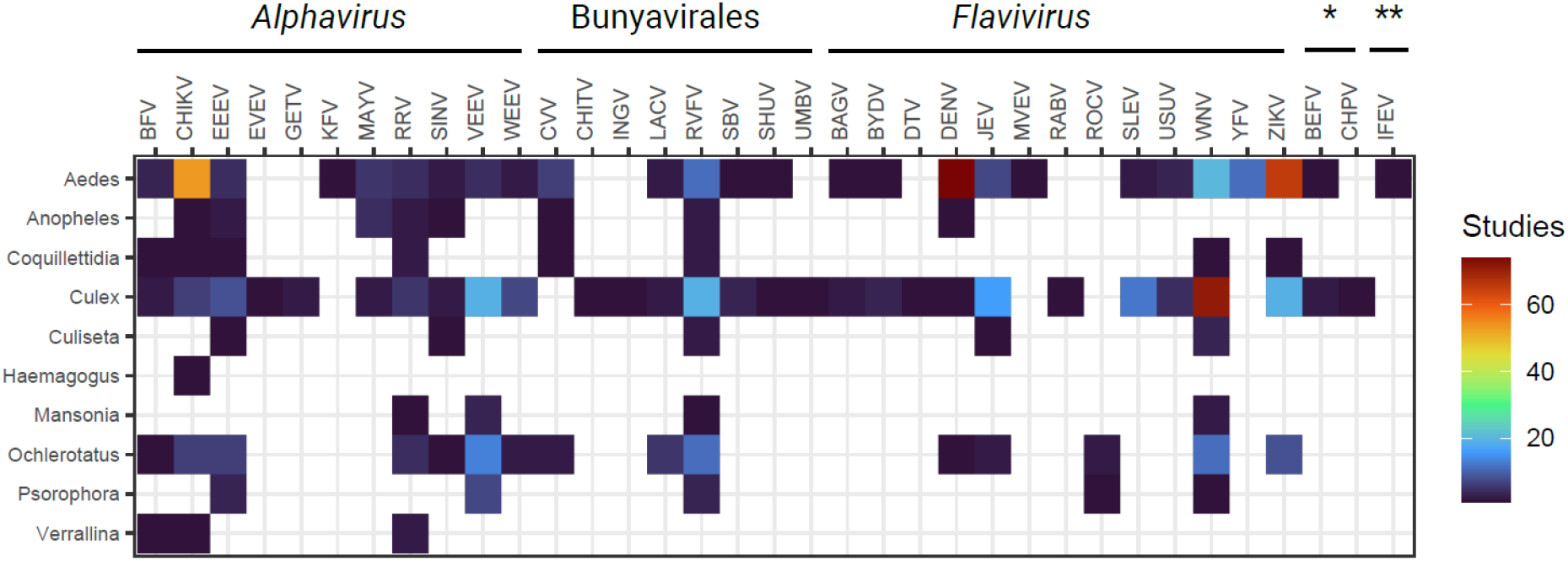
Number of studies that have experimentally tested a given arbovirus-mosquito genus pair (as in Figure 1). Abbreviations follow naming conventions in virology. (Key: *Rhabdoviridae; \*\**Orbivirus*)

Even within tested mosquito-virus combinations, effort is distributed unevenly. A small subset of viruses and their main vectors are extremely well studied: the ten most commonly tested combinations were dengue virus in *Aedes aegypti* (n = 47 studies) and *Ae. albopictus* (n = 19), Zika virus in *Ae. aegypti* (n = 35) and *Ae. albopictus* (n = 18), chikungunya virus in *Ae. aegypti* (n = 24) and *Ae. albopictus* (n = 23), and West Nile virus in *Culex pipiens* (n = 30), *Cx. quinquefasciatus* (n = 16), and *Cx. tarsalis* (n = 8). Most research has focused on flaviviruses (Flaviviridae: *Flavivirus;* n = 13 viruses, 180 studies) and alphaviruses (Togaviridae: *Alphavirus;* n = 11 viruses, 79 studies), with less focus on bunyaviruses (Bunyavirales; n = 8 viruses, 26 studies), rhabdoviruses (Rhabdoviridae; n = 2 viruses, 2 studies), and orbiviruses (one study on Ife virus). Even within virus species, distribution of effort was often unequal: for example, dengue serotype 2 (DENV-2), the most readily available lineage for experimental work, is far better studied (n = 90 studies) than the other three serotypes (DENV-1: n = 30, DENV-3: n = 17, DENV-4: n = 13). Focus on particular viruses over time tended to track the timing of major outbreaks (Figure 3), including the emergence of West Nile virus (1999-), chikungunya (2013-2014), and Zika virus (2015-2016) in the Americas.

**Figure 3.**
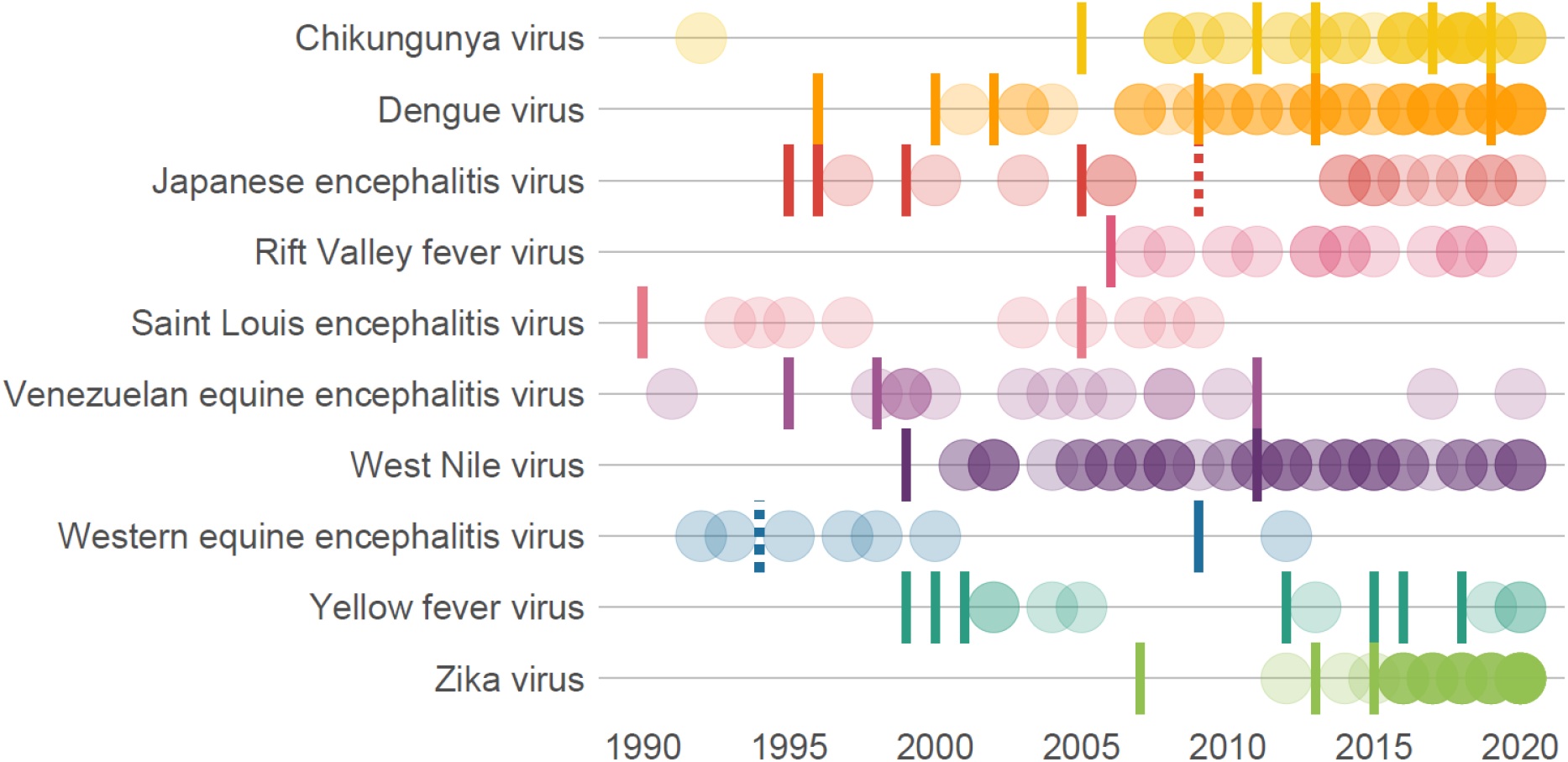
Number of studies over time, broken down by the 10 viruses that appeared in the most studies. (Opacity is proportional to the number of studies in any given year.) Tick marks represent notable outbreaks that motivated further inquiry, broken down by pathogen, and sourced from primary literature, including the WHO Disease Outbreak News.^1^ In some cases, outbreaks for one disease may have increased interest in others, e.g., CHIKV in India in 2005, and ZIKV virus in Yap in 2007, were followed by substantial renewed interests in dengue; the emergence of CHIKV and ZIKV in the Americas in 2013-15 appear to have been accompanied by incidental research on Japanese encephalitis virus, likely in many of the same experiments.

The viruses studied in the widest range of mosquitoes were not always the most-studied viruses overall; the ten viruses studied in the widest range of mosquito species were Rift Valley fever virus (RVFV; n = 36 species), West Nile virus (WNV; n = 36), Venezuelan equine encephalitis virus (VEEV; n =31), Zika virus (ZIKV; n = 26), Eastern equine encephalitis virus (EEEV; n = 24), chikungunya virus (CHIKV; n = 21), Ross River virus (RRV; n = 19), dengue virus (DENV; n = 13), Japanese encephalitis virus (JEV; n = 13), and Barmah Forest virus (BFV; n = 8).

### Geographic coverage

Lineage variation in vector competence can be quite striking, even within the geographic range of a single globalized species like *Ae. aegypti* or *Ae. albopictus* (Vega-Rúa et al., 2014). To explore how arboviral research captures this dimension of natural variation, we recorded where each study’s mosquitoes were sourced from. The majority of studies have used mosquitoes sourced from the United States (Figure 4), followed distantly by Brazil and Australia. The range of pathogens tested using mosquitoes from a given country is slightly more even, but still, the majority of work has focused on the United States, Brazil, Australia, China, India, and western Europe (Figure 5). By both measures, Africa and eastern Europe have been severely understudied, particularly compared to the Americas, where multiple explosive multinational epidemics have forced researchers to answer questions about broad geographic risk.

**Figure 4.**
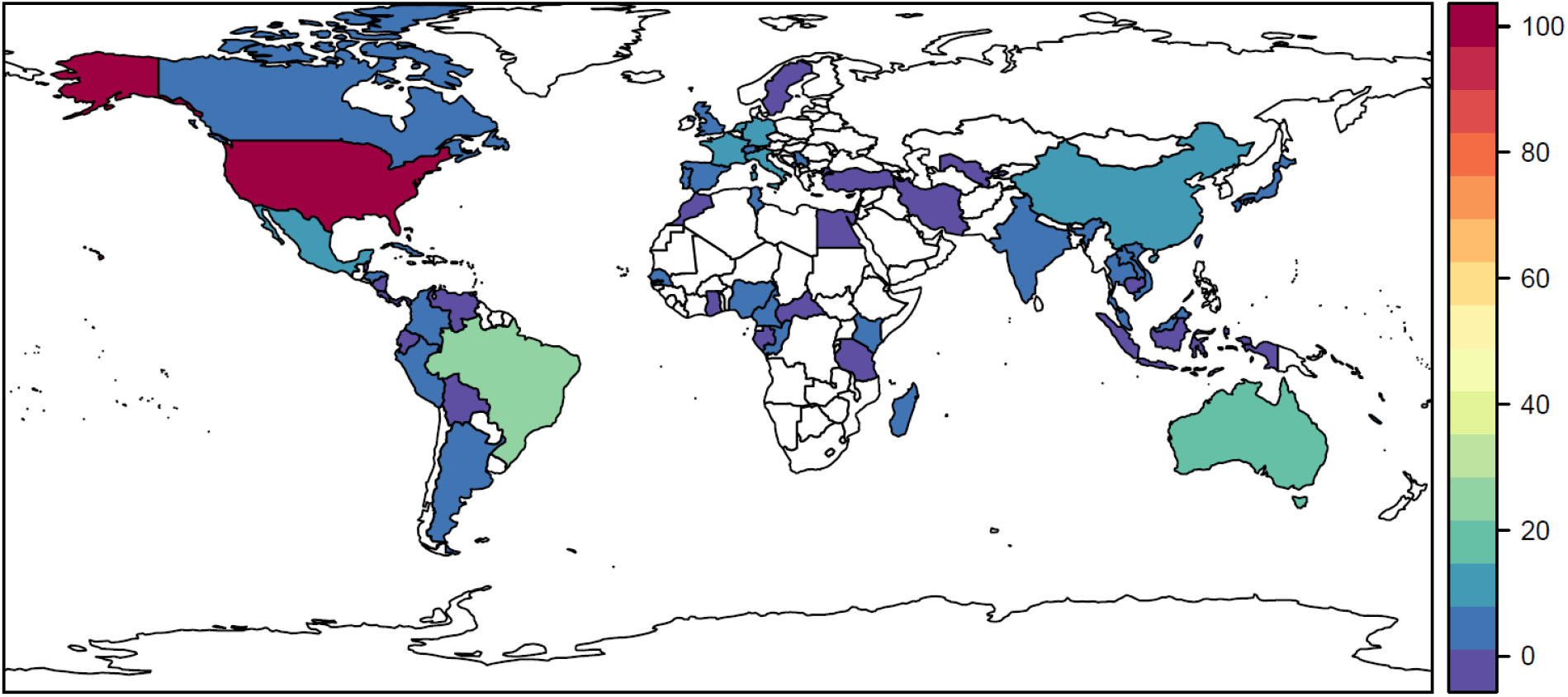
Number of studies that use mosquitoes sourced from a given country.

**Figure 5.**
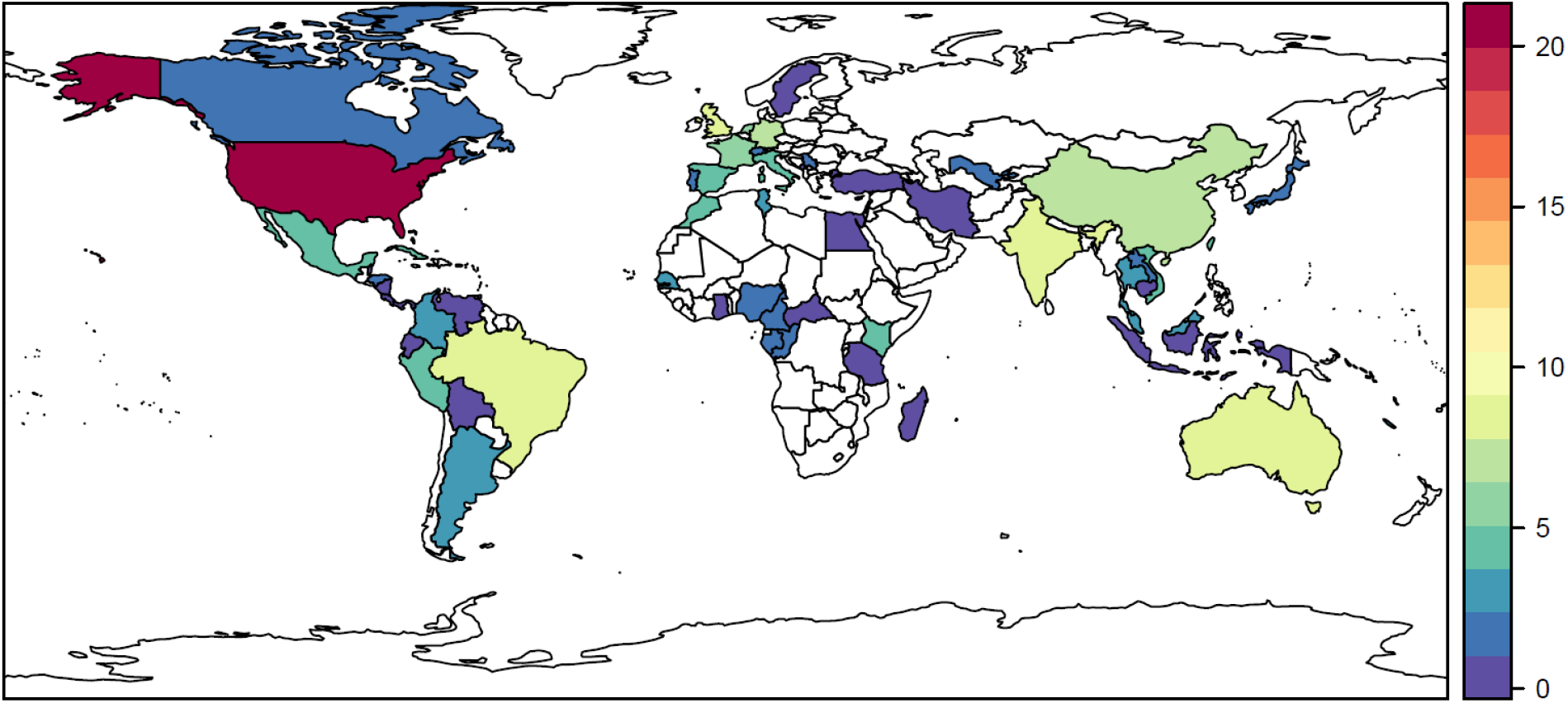
The number of pathogens (out of a total 35 distinct viruses recorded in our review) that have been tested using mosquitoes sourced from a given country.

## Discussion

In total, we identified 265 studies in Web of Science that reported the results of vector competence experiments with mosquitoes and their arboviruses. Our results suggest that research effort in this field has largely been driven by the shifting priorities of arboviral outbreak response. As such, there are numerous gaps in viral taxonomy, vector taxonomy, and vector geography that limit the utility of available data for future outbreak response. Moreover, the vast majority of mosquito-virus pairs (> 90%) are simply untested within our sample of studies, indicating that network-level understanding of arbovirus ecology is incomplete. In particular, any possibly-observable network is constrained to a very narrow subset of possible combinations (Figure 6), which may undermine efforts to accurately model the mosquito-arbovirus network (Dallas et al., 2021; Evans et al., 2017). Some gaps may particularly limit ecological inference: for example, anthropophilic vectors like *Ae. aegypti* are far better studied than bridge and sylvatic vectors like *Ae. africanus, Sabethes* spp., or *Haemagogus spp*. These species may be less important during epidemics, but determine the risk of an enzootic virus reaching humans and, conversely, whether or not sylvatic cycles are established after an epidemic ends (Althouse et al., 2016; Dallas et al., 2021).

**Figure 6.**
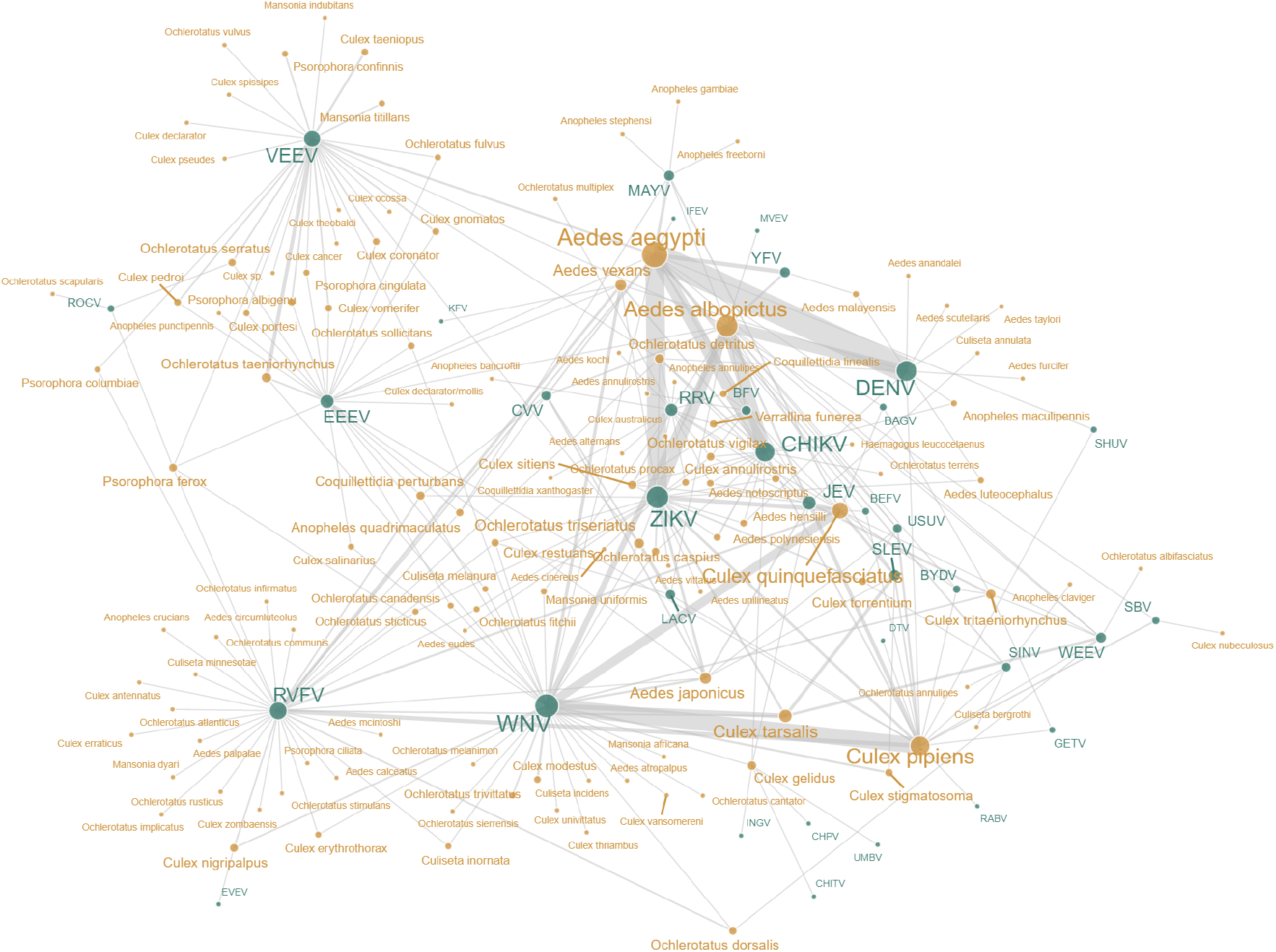
Network visualization of arbovirus (green) and mosquito (yellow) pairs experimentally tested. Notably, any observable network of compatible species would be constrained to “fit” inside this sampling-based network, highlighting how experimental effort determines observable pairs of compatible species more than their biology. Abbreviations follow naming conventions in virology. Nodes represent virus and mosquito species, where node size is proportional to the number of studies involving each (range for mosquitos: 1-136; range for viruses: 1-109). Edges represent a record of an experimentally tested pair, where edge weight is proportion to number of studies returned for each pair (range: 1-47). Edges do *not* record species compatibility.

Efforts to fill these gaps could take several approaches. First, researchers can use network science to identify important vector-virus pairs and conduct vector competence experiments that would fill key knowledge gaps. Just as machine learning can be used to guide species sampling for viral discovery in nature (Becker et al., 2022), predictive models can be used to develop or augment shortlists of the vectors that will be most relevant during emergency scenarios that have previously been flagged (e.g., yellow fever establishment in the Asia-Pacific region (Wasserman et al., 2016)). The benefits of model-experiment feedbacks are iterative: at present, many models will struggle with the sparsity of the vector-virus matrix, but targeting important sources of uncertainty will lead to better predictions.

Second, researchers can work towards a more cohesive geographic picture of how risk varies, both for cosmopolitan vectors like *Ae. aegypti* and *Ae. albopictus*, and for vectors that are locally abundant or are known to feed on amplifying hosts (Hartman et al., 2019). The geographic biases we describe here reflect where a relatively small number of institutions are able to continuously fund and maintain vector colonies. This process is itself often extractive and inefficient in nature (i.e. mosquito colonies established with species from countries facing public health emergencies are often used by researchers in the United States and Europe to generate high-impact publications). Deeply linked to capacity and sustainability of institutions housing laboratory colonies, a lack of training in medical entomology has been recently noted as a global issue in the response to outbreaks of vector-borne diseases (APMEN VCWG, 2020; Connelly, 2019; Sifferlin, 2018).

Finally, our study highlights that a substantial breadth and depth of vector competence data are published every year in the peer-reviewed literature, but currently, the results of these experiments have no standardized home. Other recent studies highlight that synthesis of these data is possible, despite the complexity of metadata required to describe variation in experimental protocols (Kain et al., 2022); however, our study highlights the challenges of recovering “findable” data from the vector competence literature. Limitations we encountered included: (1) several databases capture these studies (and older studies in particular appear to be missing from Web of Science); (2) keywords we used may not have captured all of the relevant studies, due to variable terminology; (3) a handful of non-English language publications—in particular, Spanish and French language publications from Latin America and Africa, respectively—will not have been captured by our search terms; and (4) not all studies reported reusable experimental results and metadata. Developing a standardized database of vector competence experiments with direct user contributions—ideally supported by a universal set of minimum data and metadata standards (Wu et al., 2022)—would help translate the inconsistent and patchy funding in this field into more immortalized data. In doing so, such a database would help researchers identify geographic and taxonomic knowledge gaps further in advance of public health emergencies, and would make more data immediately available to public health agencies in a searchable format once an outbreak begins.

As we show here, data science approaches can be useful to identify trends and gaps in scientific understanding of arboviral ecology and evolution (Albery et al., 2021). In some cases, filling those gaps will be more challenging: for example, establishing colonies is harder for some mosquito species than others, and some viruses require higher biosafety levels, limiting the number of researchers with the ability to safely work with them. Future work may also aim to expand our scope to other medically important vectors, including ticks and midges (for which we excluded a handful of studies that were identified by our search terms). Similarly, future work could examine trends in how coinfection dynamics are studied and tested; both insect-specific viruses and *Wolbachia* have been considered as potential biological countermeasures to arboviral transmission in mosquitoes, but the network of pathogen-coinfection-vector combinations has been characterized even less systematically, and these data remain largely unsynthesized. Addressing these types of questions in the future may point to new opportunities for both empirical research and modeling that harnesses these data to predict and prevent arboviral emergence.

## Materials and Methods

A systematic literature search was conducted on Web of Science to identify suitable records that described vector competence experiments with mosquito-borne viruses. Our search used the following terms: “(“vector competence” OR “extrinsic incubation period” OR “vectorial capacity” OR “dissemination”) AND (arbovirus OR virus) AND (experiment* OR trial OR captive* OR laboratory) AND mosquito.” Our search returned 570 records in February 2021.

We performed an initial screen of records based on abstracts excluding reviews, methodology descriptions, and studies with no experimental infection (N = 135), studies with non-virus infections, insect-specific infections, infection regimes with confounding treatments (i.e., coinfection with *Wolbachia* or insect-specific viruses; N = 71), studies involving infection in non-mosquitoes, experimental vector infection without any reported results describing competence quantitatively, or *ex vivo* data (N = 55), and studies fitting multiple exclusion criteria (N = 22). For the remaining 287 records, we undertook a second round of screening to determine suitability (i.e., did the study include some sort of experimental test of mosquito competence for arbovirus transmission). From the 265 studies that were within scope, we recorded the species pairs of mosquitoes and arthropod-borne viruses that were experimentally tested together; the country from which wild mosquitoes were originally collected (even if populations were maintained in laboratory settings long-term); and, when available, mosquito subspecies and (as applicable) dengue virus serotype. Using the ‘taxize’ R package, we updated mosquito binomial names if a more recent valid name could be found in the NCBI taxonomic database; in all other cases, we used taxonomy reported verbatim from source studies, including non-standard naming conventions (e.g., “*Culex* sp.”, “*Culex declarator/mollis*”).

All analyses were conducted, and figures generated, in R software version 4.0.3.

## Acknowledgements

This work was supported by funding to the Viral Emergence Research Initiative (VERENA; viralemergence.org), including NSF BII 2021909. SJR was additionally supported by NSF DBI 2016265 CIBR: VectorByte: A Global Informatics Platform for studying the Ecology of Vector-Borne Diseases

## Author Contributions

BC, ARS, CJC, and SJR generated the data, and CJC and ARS generated the visualizations. All authors contributed to the conceptualization of the study and the writing of the manuscript.

## Data and Code Availability

Raw data and code to reproduce analyses are available on Github at github.com/viralemergence/bolide.

## Footnotes

1 CHIKV: 2005 – India; 2011 – New Caledonia; 2013 – the Americas; 2017 – Italy; 2019 – Republic of the Congo. DENV: 1996 – the Americas; 2000 – DENV-3 introduced to Brazil; 2002 – Brazil; 2009 – the Americas; 2013 – southeast Asia; 2019 – the Americas. JEV: 1995 – Australia; 1996 – Nepal; 1999 – India; 2005 – India. RFV: 2006 – East Africa; 2009 – WHO WPRO and SEARO reference labs created. SLEV: 1990 – Florida, USA; 2005 – Argentina. VEEV: 1995 – Colombia and Venezuela; 1998 – Colombia; 2011 – Colombia and Venezuela. WNV: 1999 – United States; 2011 – Europe and, unrelated, Australia (Kunjin virus). WEEV: 1994 – last human case in North America (dotted line); 2009 – Uruguay. YFV: 1999 — South America and importation into United States; 2000 – Guinea; 2001 – Côte d’Ivoire; 2005 – Sudan; 2012 – Sudan; 2015 – Angola and DRC; 2016 – Brazil; 2018 – Brazil. ZIKV: 2007 - Yap Island; 2013 – French Polynesia; 2015 – the Americas. (Azar et al., 2020; Carlson et al., 2022; de Araújo et al., 2012; Diaz et al., 2018; Gutiérrez-Bugallo et al., 2019; Guzmán-Terán et al., 2020; Klitting et al., 2018; Meganck and Baric, 2021; Musso et al., 2018; Nathan et al., 2001; Xu et al., 2022)

